# The spatial spread and the persistence of gene drives are affected by demographic feedbacks, density dependence and Allee effects

**DOI:** 10.1101/2024.08.08.607064

**Authors:** Léna Kläy, Léo Girardin, Vincent Calvez, Florence Débarre

## Abstract

Homing gene drive alleles bias their own transmission by converting wild-type alleles into drive alleles. If introduced in a natural population, they might fix within a relatively small number of generations, even if they are deleterious. No engineered homing gene drive organisms have been released in the wild so far, and modelling is essential to develop a clear understanding of the potential outcomes of such releases. We use deterministic models to investigate how different demographic features affect the spatial spread of a gene drive. Building on previous work, we first consider the effect of the intrinsic population growth rate on drive spread. We confirm that including demographic dynamics can change outcomes compared to a model ignoring changes in population sizes, because changes in population density can oppose the spatial spread of a drive. Secondly, we study the consequences of including an Allee effect, and find that it makes a population more prone to eradication following drive spread. Finally, we investigate the effects of the fitness component on which density dependence operates (fecundity or survival), and find that it affects the speed of drive invasion in space, and can accentuate the consequences of an Allee effect. These results confirm the importance of checking the robustness of model outcomes to changes in the underlying assumptions, especially if models are to be used for gene drive risk assessment.

## 1 Introduction

A promising but controversial new strategy for the control of natural populations, artificial gene drive biases the transmission of particular alleles to the offspring, over expectations of regular Mendelian transmission [1–3]. Such alleles can be detrimental to the individuals carrying them, and yet spread in a population thanks to their transmission advantage. Artificial gene drive implementations, so far still restricted to laboratory settings, have achieved transmission rates of 99% in yeast (*Saccharomyces cerevisiae*, [4]), more than 90% in mosquitoes (*Anopheles gambiae*, [5]), and more than 85% in fruit flies (*Drosophila melanogaster*, [6]).

In “homing drives”, biased inheritance relies on gene conversion: in a heterozygous cell, the gene drive cassette present on one chromosome induces a double-strand break on a target site on the homologous chromosome, and repair by homologous recombination duplicates the cassette. The repetition of this process through generations favors the propagation of the drive allele in the population. Conversion can theoretically happen at different steps of the life-cycle, like in the germline of the parents, or in the zygote. Practical implementations in the lab have focused on conversion in the germline [7].

Biased transmission via gene conversion can lead to the spread of new, potentially deleterious traits in a population within a relatively small number of generations. Two main types of drive can be distinguished: *replacement drives*, aiming to change features of the target population without directly affecting its size, and *suppression drives*, aiming to reduce population size (an extreme being *eradication drives*). Because we are interested in exploring the effect of demographic dynamics on the spatial spread of gene drive alleles, our work here will focus on suppression drives. Experimental proofs of principle for this type of drive have been obtained with cage populations [8, 9], and the feasibility in large populations has been confirmed by theoretical studies [2, 10, 11].

Artificial gene drive like CRISPR-based homing drive holds promise for addressing a number of important real-world issues [12–14], among which the burden caused by vector-borne diseases like malaria. Artificial gene drive could be used to spread a new trait rendering progeny of vector mosquitoes unable to transmit disease [15], or simply leading to the reduction of vector mosquitoes population size over time [8, 16]. Applications of artificial gene drive are however not limited to human health. Gene drive could help conserve or even partially restore native ecosystems by disadvantaging invasive species or favouring endemic ones [17, 18]. It could also be used in agriculture to reverse insecticide resistance in pest animal species [19] or make weeds susceptible again to herbicides [20].

As of today, no artificial gene drive organisms have been released in the wild. Lab experiments, as well as mathematical and computational models, are crucial to evaluate the risks and benefits of gene drive, and to assess the safety of potential releases. Models are however simplifications of the living world, and it is crucial to understand the impact and importance of various modelling choices, and to test the robustness of results to changes in modelling assumptions.

The simplest theoretical models of gene drive represent well-mixed populations [21], and focus on allele frequencies changes over time [18, 22–25]. Here, we investigate the spatial spread of a gene drive allele, and how demographic features affect it. Previous work has shown that propagation of a drive in a well-mixed population did not necessarily imply that the drive would spread spatially. This is in particular the case when the drive is threshold-dependent, i.e. when, in a well-mixed population, it needs to be introduced in a high enough amount to increase in proportion [26, 27]. While changes in population density may be ignored when a drive barely affects reproduction or survival, it becomes important to consider them in the case of a suppression drive, because its increase in proportion directly affects population size. Previous work on a specific model [11, 28] found that demographic features can affect the speed of advance of a drive wave over a continuous space. Here, we will assess the robustness of this result to different modelling assumptions.

A population’s growth rate is determined by birth and death rates [29]. Density regulation may affect the two differently, which has consequences for overall demographic dynamics [30]. Likewise, which fitness component is affected by the drive (i.e., whether the drive reduces fecundity or decreases survival) can also influence outcomes [31]. Finally, growth at low population density may be different from growth at high population densities, i.e. Allee effects may operate [32]. This can be caused by inbreeding depression, or difficulties to find a mate when the population density is low, for example [33]. Allee effects are frequently observed in the wild, including for animals considered as potential targets of control by artificial gene drive, like mosquito species affected by inbreeding depression [34–36]. The existence of Allee effects may also influence the outcome of the release of a drive affecting population size.

In this article, we consider a one-dimensional continuous environment, and we study the spatial spread (or failure thereof) of a drive allele invading an established wild-type population. We follow the densities of the different genotypes (drive homozygous, wild-type homozygous and heterozygous) over space and time using partial differential equations. We compare four types of demographic models, depending on the presence or absence of an Allee effect, and the fitness component (birth or death) on which density dependence operates. We characterise the spread of a drive by the existence and direction of its wave of advance, by the final total population density after the drive has spread (or failed to), and by the speed of the wave. We find that an Allee effect might help to eradicate or reduce the density of the targeted population, but that it might also lead to the failure of threshold-dependent drive invasions. We also find that the effect of demography on drive spread is limited in the case of density regulation on the birth rate, but is not when density regulation affects the death rate, where wave speed increases with intrinsic growth rate. This difference emerges because drive invasion over space primarily relies on the birth of new individuals. These results highlight the importance of ecological details on the outcome of the release of a drive.

## 2 Models and methods

### 2.1 Models

In this section, we build step-by-step the different models that we will compare. These models differ in their demographic components, which we first introduce.

#### 2.1.1 Demographic terms

To assess how sensitive results might be to different demographic modelling choices, we will consider four models differing in their birth and death terms. We first illustrate these four demographic models in the case of a genetically and spatially homogeneous population, composed only of wild-type individuals. We will compare density dependence acting on the birth term (Models ℬ𝒩 and ℬ𝒜) or death term (Models 𝒟𝒩 and 𝒟𝒜), and the absence (Models ℬ𝒜 and 𝒟𝒜) and the presence of an Allee effect (Models and).

We denote by *r* the population’s intrinsic growth rate, and by *a* the parameter controlling the Allee effect threshold (when there is an Allee effect, *™*1 *≤ a ≤* 1). In these models, the population’s initial growth rate (i.e., when *n →* 0) is *r* in the absence of Allee effect, and *™a r* in the presence of Allee effect. When *™*1 *< a <* 0, the Allee effect is said to be weak (the initial growth rate remains positive), while when 0 *< a <* 1, the Allee effect is said to be strong (the initial growth rate is negative; the population only grows if already at high enough density; see Appendix B for details).

Population density is scaled so that the carrying capacity in all models is 1, and time is scaled so that the death rate in the absence of density regulation is 1. Denoting by *n* (*t*) population density at time *t*, the four models read, before introducing genetic diversity and spatial variation, as follows:

Model ℬ𝒩 (Density regulation on birth terms; no Allee effect)

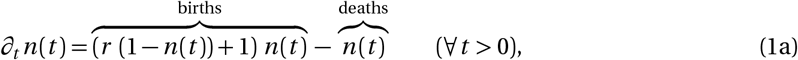

Model ℬ𝒜 (Density regulation on birth terms; Allee effect present)

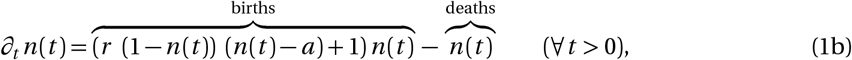

Model 𝒟𝒩 (Density regulation on death terms; no Allee effect)

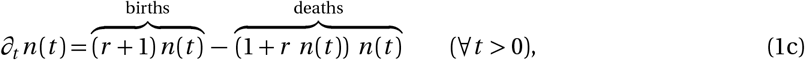

Model 𝒟𝒜 (Density regulation on death terms; Allee effect present)

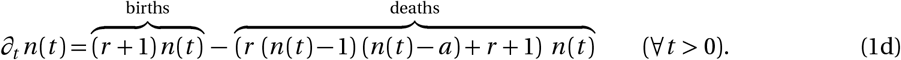

In the following, we denote the birth rate by *B* (*n* (*t*)) and the death rate by *D* (*n* (*t*)) such that the four equations can all be written as:

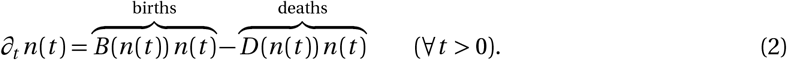

#### 2.1.2 Drive and wild-type

The demographic models being defined, we now add genetic diversity to the models, following the same approach as in [11, 28]. The variable *n* (*t*) becomes the total density, and we denote by *n*_*i*_ the density of individuals with genotype *i*. There are two possible alleles at the locus that we consider: the wild-type allele (W) and the drive allele (D), so that there are three different genotypes: wild-type homozygotes (*i* = WW), drive homozygotes (*i* = DD) and heterozygotes (*i* = DW). The fitness effect of a genotype is represented by a coefficient *f*_*i*_ acting on the birth term. It represents the selective disadvantage conferred by the drive to the individual carrying it. Wild-type homozygotes have fitness *f*_WW_ = 1, drive homozygotes have fitness *f*_DD_ = 1 *™ s*, where *s* is the fitness cost of the drive, and drive heterozygotes have fitness *f* = 1 *™ sh*, where *h* is the dominance parameter. We assume that mating occurs at random: the probability that a genotype *l* mates with a genotype *k* is equal to 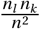. Finally, we denote by 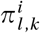 the probability for a couple of parents with genotypes *l* and *k* to have offspring of type *i*. This probability depends on the moment at which gene conversion takes place and on the probability *c* that gene conversion takes place and is successful (0 *≤ c ≤* 1). Here we assume that gene conversion takes place in the germline, because this is the timing currently successfully implemented in the lab [7, 37], unlike gene conversion in the zygote. We assume that failed gene conversion leaves the wild-type allele intact. In other words, we do not consider here the generation of resistance alleles. With these assumptions, the dynamics are now given by the following equations:

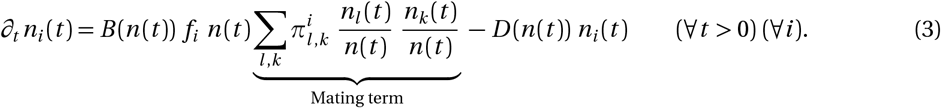

The formulas for 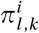 are included in the full equations presented in Appendix C.1.

#### 2.1.3 Space

Our equations so far did not include space; we now add this component. We assume that the movement of individuals is described by a diffusion term with equal diffusion coefficients for all genotypes. Space is scaled such that these coefficients are normalised to 1. We obtain the following equations:

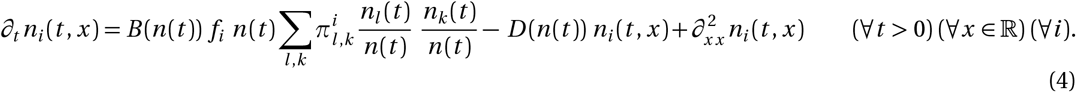

All parameters of the models are summarised in Table 1.

**Table 1:** Model parameters.

| Parameters | Range values | Description |
| --- | --- | --- |
| $r$ | $(0, +\infty)$ | Intrinsic growth rate |
| $c$ | $[0, 1]$ | Conversion rate |
| $s$ | $(0, 1)$ | Fitness cost of drive homozygotes |
| $h$ | $[0, 1]$ | Drive dominance |
| $a$ | $[-1, 1]$ | Allee effect threshold |

We have presented equations with genotype densities *n*_*i*_ (condensed model in equation (4); full equations for each model are given in Appendix C). The model can be rewritten to follow allele densities instead (see Appendix C.2), or total population size and allele frequencies (see Appendix C.3); different steps of the analysis may require different formulations of the model.

### 2.2 Traveling waves

The introduction of drive individuals in a wild-type population will give rise to a wave of change in genotype densities through space, called a traveling wave (except in the *gene drive clearance* case, see below). Traveling waves propagate with a constant speed, while maintaining their shape in space. We consider an initial condition in which the left half of the domain is full of drive (*n*_DD_ = 1), and the right half is full of wild-type (*n*_WW_ = 1), as illustrated in Figure S1. In this article, we are not exploring the effect of inoculum size and distribution, which is a question in itself, and arises in particular in the case of threshold-dependent drives [38]. We therefore choose an initial condition maximizing the possibility of drive spread. The model is then solved numerically. We classify the outcomes into five categories, present in the four models, depending on: the existence or not of a traveling wave; whether the population persists or is eradicated; and in the former case, the genotype(s) present at the end. These outcomes are illustrated in Figure 1.

**Figure 1:**
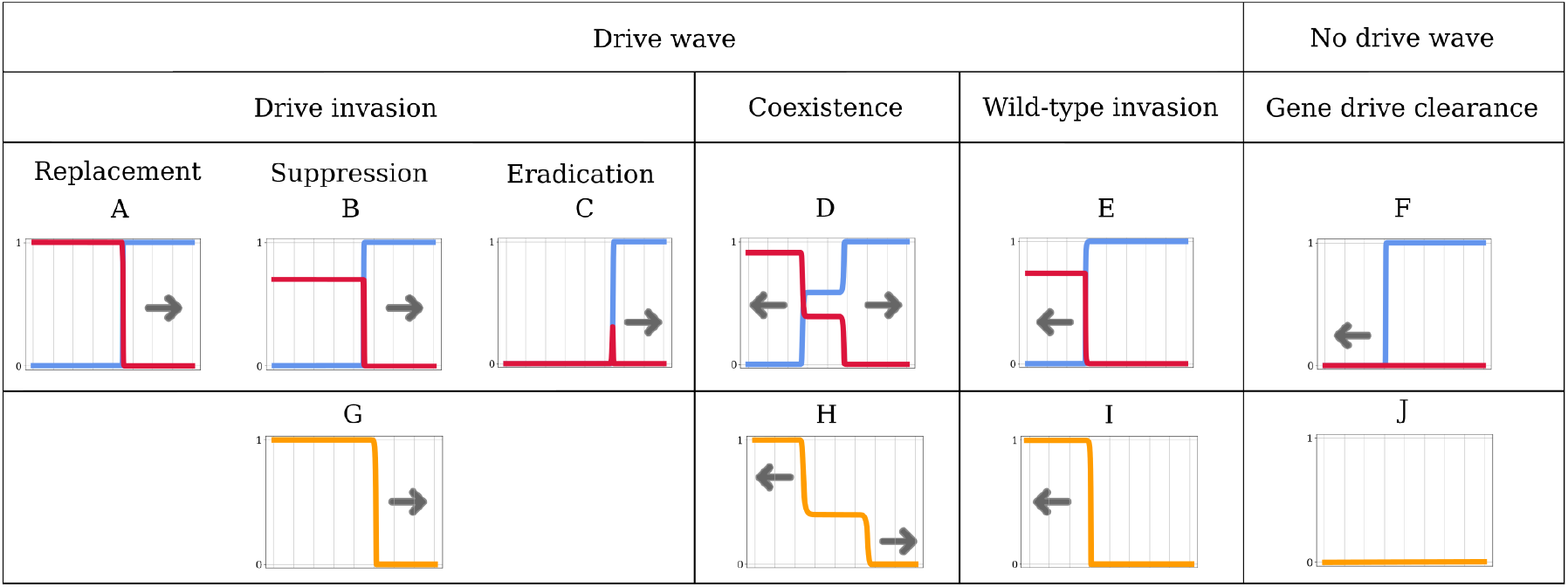
Types of spatial dynamics. Panels A–F correspond to allele densities, with the drive allele in red and the wild-type allele in blue. Arrows represent the direction of advance of the wave. Panels G–J show the equivalent with the drive allele frequency. The horizontal axis represents space.

It can happen that the model leads to the decay of the drive allele uniformly in space. This case arises in particular when a well-mixed population composed only of drive individuals is not sustainable. The introduced drive subpopulation just dies out, freeing space; in this case, there is no drive traveling wave. We describe this as *gene drive clearance* (Figure 1F, J). The wild-type population then recolonizes the emptied space, at a speed described in the standard Fisher-KPP traveling wave problem (see [39–41]).

When the drive traveling wave does exist, we distinguish between two cases, depending on the sign of the speed *v*. When *v >* 0, the wave moves to the right: it is a *drive invasion*. When *v <* 0, the wave moves to the left: it is a *wild-type invasion*. In some specific cases, drive and wild-type invasions can happen simultaneously: the waves decompose into two sub-traveling wave solutions over half of the domain. They move in opposite directions and lead to the coexistence of both alleles in-between (Figure 1D, H).

When the drive invades and replaces the wild-type (Figure 1G), we distinguish three cases depending on the state of the population in the wake of the front(s): i) in the case of *replacement* drives, the population persists in the wake of the front(s) at the same density as the original wild-type population (Figure 1A); ii) in the case of *suppression* drives, the population persists in the wake of the front(s), albeit at a lower density than the original wild-type population (Figure 1B); iii) in the case of *eradication* drives, the population is eradicated in the wake of the drive invasion front(s), leaving empty space behind (Figure 1C). In the latter case, two scenarios are possible: persistence of drive homozygotes only (as in Figure 1G); but also possibly persistence of all genotypes (as in Figure 1H).

The code for these simulations is available on GitHub (https://github.com/LenaKlay/gd_project_1/deterministic). We ran our simulations in Python 3.6, with the Spyder environment. Heatmaps in Figures 3 were computed thanks to the INRAE Migale bioinformatics facility.

## 3 Results

### 3.1 Demography and dominance can affect the final allelic proportions

Here, we focus on the importance of demography in the model, i.e., on the role played by the intrinsic growth rate *r* on the final allelic proportions. Analytical results can be obtained for *r* = 0 and for *r → ∞*; intermediate cases are investigated numerically.

**Table 2:** Types of model outcomes for Models ℬ𝒩, ℬ𝒜, 𝒟𝒩 and 𝒟𝒜, depending on the fitness cost *s*, intrinsic growth rate *r* and dominance parameter *h*. The outcomes are in terms of allele proportions, as in Fig. 1G–J.

| (a) When $h < 1/2$ | | | |
| --- | --- | --- | --- |
| | $0 < s < s_1$ | $s_1 < s < s_{2,g}$ | $s_{2,g} < s < 1$ |
| $r \rightarrow \infty$ | Drive invasion | Coexistence | Wild-type invasion |
| $r = 0$ | Drive invasion** | | Gene drive clearance |

**Table 2:** Types of model outcomes for Models $\mathcal{BN}$ , $\mathcal{BA}$ , $\mathcal{DN}$ and $\mathcal{DA}$ , depending on the fitness cost $s$ , intrinsic growth rate $r$ and dominance parameter $h$ . The outcomes are in terms of allele proportions, as in Fig. 1G–J.
| (b) When $h > 1/2$ | | | |
| --- | --- | --- | --- |
| | $0 < s < s_{2,g}$ | $s_{2,g} < s < s_1$ | $s_1 < s < 1$ |
| $r \rightarrow \infty$ | Drive invasion | Bistability | Wild-type invasion |
| $r = 0$ | Drive invasion** | Gene drive clearance | |

When *r* = 0, deaths and births compensate each other in a fully wild-type population. In this limit case, models ℬ𝒩, ℬ𝒜, 𝒟𝒩 and 𝒟𝒜 are the same (given in equation (C.5)). Both the final densities of all genotypes and the speed of the wave are therefore the same for all models, which we will characterise below, recalling results from our previous work [28].

Leaving aside the density-dependence constraint, the bigger *r* is, the faster the wild-type population grows. When *r → ∞*, final allelic proportions are the same in models ℬ𝒩, ℬ𝒜, 𝒟𝒩 and 𝒟𝒜 (see Section C.4). This is however not necessarily the case for the total population density and for the wave speed.

Following previous work [28], let us introduce:

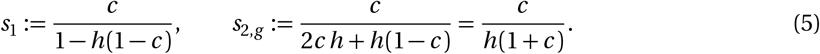

These are threshold values of the fitness cost *s* determining qualitatively different outcomes. When the drive allele is recessive (*h <* 1*/*2), *s*_1_ *< s*_2,*g*_ ; when the drive allele is dominant (*h >* 1*/*2), *s*_1_ *> s*_2,*g*_.

When the fitness cost *s* is low enough (*s <* min(*s*_1_, *s*_2,*g*_)), there is a wave of advance of the drive for both *r* = 0 and *r → ∞* (drive invasion, as in Fig. 1G).

When the fitness cost *s* is high enough (*s >* max(*s*_1_, *s*_2,*g*_)), and the intrinsic growth rate is high (*r → ∞*), the drive wave retreats (wild-type invasion, as in Fig. 1I). When the intrinsic growth rate is low (*r* = 0), *s >* max(*s*_1_, *s*_2,*g*_) results in drive clearance (as in Fig. 1J): the drive is just too costly even for a full-drive population.

What happens for intermediate fitness cost (min(*s*_1_, *s*_2,*g*_) *< s <* max(*s*_1_, *s*_2,*g*_)) and high growth rate depends on the dominance parameter *h*. If *h <* 1*/*2, drive and wild-type alleles coexist eventually (coexistence, as in Fig. 1H). If *h >* 1*/*2, there is a bistability, the drive is threshold-dependent: the final outcome is either drive invasion or wild-type invasion, and depends on the initial conditions.

These results are summarised in Table 2.

These results illustrate the importance of taking demography into account. Threshold-dependent drives (i.e. drives leading to bistabilities), are considered more socially responsible than threshold independent drives, as they are potentially localised and reversible [25, 26, 42]. The intrinsic growth rate *r* is a key component to reach this threshold dependence, as *r* has to be sufficiently large for the bistability to happen. Indeed, a small *r* would result in the systematic decay of gene drive alleles (Table 2) and no possibility of drive invasion at all.

As in models without demography nor spatial structure [22, 23], the dominance parameter *h* determines whether threshold-dependence can be attained or not: a bistable outcome only exists when 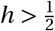, i.e. when the fitness of heterozygous individuals is closer to the fitness of drive homozygous individuals than to that of wild-type homozygous individuals. This result was already given in a simpler panmictic model [22]: indeed, the birth and death terms in our models ℬ𝒩 and ℬ𝒜 tend to this panmictic model for large values of *r*.

### 3.2 The strength of the Allee effect and the choice of the fitness component targeted by the density-dependence affect the final allelic density

In the previous section, we have only described outcomes in terms of allele frequencies. In this section, we compare the final population density *n*^*^ in the four models, and in particular conditions for which the population goes extinct (*n*^*^ = 0). We detail the final densities in all three types of invasions: drive invasion, wild-type invasion, and coexistence. In case of gene drive clearance (decay of the drive allele uniformly in space), the final density is equivalent to the one obtained after a wild-type invasion: population size goes back to carrying capacity 1).

In all three types of invasion, there are up to three possible regimes: population eradication (*n*^*^ = 0); population persistence (*n*^*^ = *n* ^+^ *>* 0); and bistability (the final total population size is either 0 or *n* ^+^ depending on the initial condition relative to a specific density *n*^*τ*^). Note that the bistability involved here is different from the bistability on allele frequencies as seen in the previous section; the bistability that we consider in this section is about population densities.

We can write the final population densities in a generic manner for the three types of invasion. We define the mean fitness ℱ:

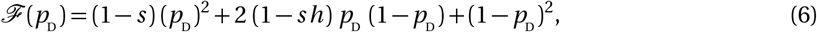

and 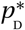 the final proportion of the drive allele in the population. It verifies:

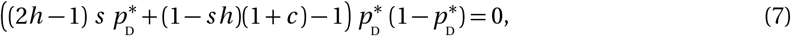

according to the allelic frequency systems detailed in Section C.3. For wild-type invasion, 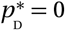 and 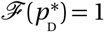; for drive invasion, 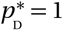 and 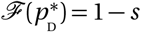; for coexistence,

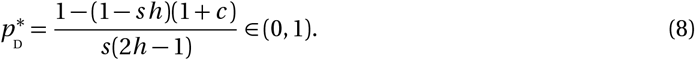

and 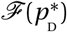 is still given by equation (6). Note that 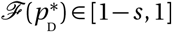 as ℱ is a decreasing function of 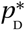.

The final densities *n*^*^ are then computed by solving the allelic frequency systems detailed in Section C.3 with the relevant value of 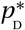. These results, holding for all values of the intrinsic growth rate *r*, are summarised in Table 3 and illustrated in Figure 2 with *c* = 0.85 and *h* = 0.9.

**Table 3:** Regimes and final densities in Models ℬ𝒩, ℬ𝒜, 𝒟 𝒟𝒩 and 𝒟𝒜, where 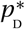 is the final proportion of the drive allele in the population and 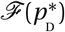 the mean fitness. We consider three regimes regarding the value of the final population density, *n*^*^: population eradication (*n*^*^ = 0), population persistence (*n*^*^ = *n* ^+^ *>* 0) and bistability (the final total population size is either 0 or *n* ^+^ depending on the initial condition relative to the threshold density *n*^*τ*^).

| Model | Regime | $n^+$ and $n^\tau$ (if it exists) |
| --- | --- | --- |
| $\mathcal{BN}$ | Eradication if $r < \frac{1-\mathcal{F}(p_D^*)}{\mathcal{F}(p_D^*)}$<br>Persistence if $r > \frac{1-\mathcal{F}(p_D^*)}{\mathcal{F}(p_D^*)}$ | $n^{+(\mathcal{BN})} = 1 - \frac{1-\mathcal{F}(p_D^*)}{r\mathcal{F}(p_D^*)}$ |
| $\mathcal{BA}$ | Eradication if $r < \frac{1-\mathcal{F}(p_D^*)}{\left(\frac{1-a}{2}\right)^2 \mathcal{F}(p_D^*)}$<br>Bistability if $r > \frac{1-\mathcal{F}(p_D^*)}{\left(\frac{1-a}{2}\right)^2 \mathcal{F}(p_D^*)}$ and $-ar < \frac{1-\mathcal{F}(p_D^*)}{\mathcal{F}(p_D^*)}$<br>Persistence if $r > \frac{1-\mathcal{F}(p_D^*)}{\left(\frac{1-a}{2}\right)^2 \mathcal{F}(p_D^*)}$ and $-ar > \frac{1-\mathcal{F}(p_D^*)}{\mathcal{F}(p_D^*)}$ | $n^{\tau(\mathcal{BA})} = \frac{1+a-\sqrt{(1+a)^2-4(a+\frac{1-\mathcal{F}(p_D^*)}{r\mathcal{F}(p_D^*)})}}{2}$<br>$n^{+(\mathcal{BA})} = \frac{1+a+\sqrt{(1+a)^2-4(a+\frac{1-\mathcal{F}(p_D^*)}{r\mathcal{F}(p_D^*)})}}{2}$ |
| $\mathcal{DN}$ | Eradication if $r < \frac{1-\mathcal{F}(p_D^*)}{\mathcal{F}(p_D^*)}$<br>Persistence if $r > \frac{1-\mathcal{F}(p_D^*)}{\mathcal{F}(p_D^*)}$ | $n^{+(\mathcal{DN})} = 1 - \frac{(r+1)(1-\mathcal{F}(p_D^*))}{r}$ |
| $\mathcal{DA}$ | Eradication if $r\left[\left(\frac{1-a}{2}\right)^2 - (1-\mathcal{F}(p_D^*))\right] < 1-\mathcal{F}(p_D^*)$<br>Bistability if $r\left[\left(\frac{1-a}{2}\right)^2 - (1-\mathcal{F}(p_D^*))\right] > 1-\mathcal{F}(p_D^*)$<br>and $r[-a - (1-\mathcal{F}(p_D^*))] < 1-\mathcal{F}(p_D^*)$<br>Persistence if $r\left[\left(\frac{1-a}{2}\right)^2 - (1-\mathcal{F}(p_D^*))\right] > 1-\mathcal{F}(p_D^*)$<br>and $r[-a - (1-\mathcal{F}(p_D^*))] > 1-\mathcal{F}(p_D^*)$ | $n^{\tau(\mathcal{DA})} = \frac{1+a-\sqrt{(1+a)^2-4(a+\frac{(r+1)(1-\mathcal{F}(p_D^*))}{r})}}{2}$<br>$n^{+(\mathcal{DA})} = \frac{1+a+\sqrt{(1+a)^2-4(a+\frac{(r+1)(1-\mathcal{F}(p_D^*))}{r})}}{2}$ |

**Figure 2:**
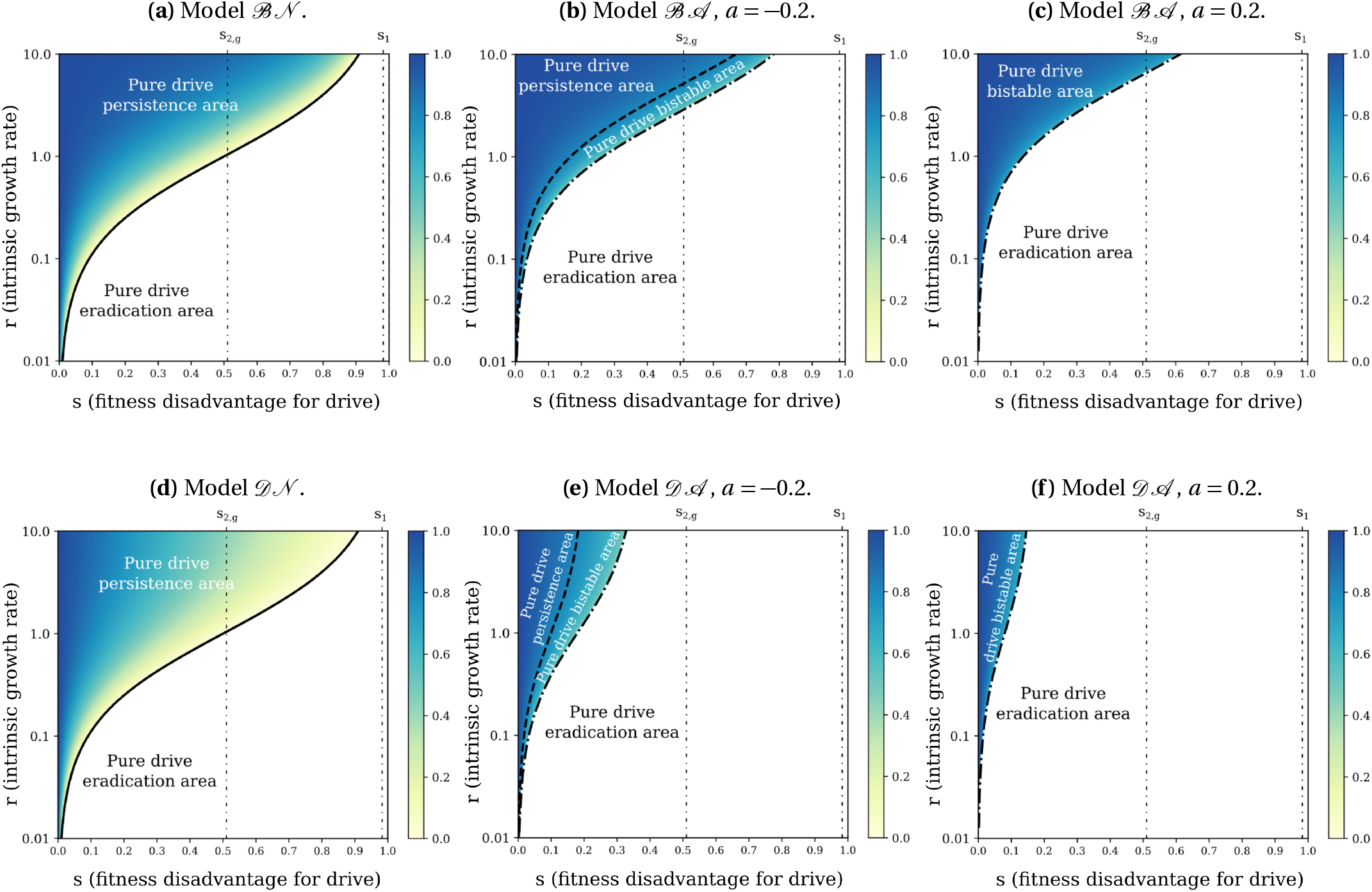
Value of the final population density *n*^*^ = max(0, *n* ^+^) in case of a drive invasion, shown in shades of color, with parameters *c* = 0.85 and *h* = 0.9. Since 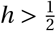, drive invasion always occurs for *s < s*_2,*g*_, is not systematic for *s*_2,*g*_ *< s < s*_1_, and never occurs for *s*_1_ *< s* (see Table 2b). The “pure drive” areas correspond to the final population density expected in case of a drive invasion: persistence (*n*^*^ = *n* ^+^), eradication (*n*^*^ = 0) or bistability (either *n*^*^ = *n* ^+^ or *n*^*^ = 0 depending on the initial condition). These final densities are given in Table 3 with 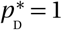, because the proportion of drive alleles in the final population is one in case of a drive invasion without coexistence.

#### 3.2.1 When the Allee effect gets stronger, the population is more prone to eradication

In the models without Allee effects, there is no bistability for the final population size once the type of invasion is known (see Table 3). However, and as seen before, the type of invasion itself might change depending on the initial conditions (bistability on allele frequency, see Table 2b), and consequently can still affect the final population size. The situation is different in models with Allee effect, where the final population size might depend on the initial conditions. In case of a weak Allee effect (*™*1 *< a <* 0), the three possible regimes are eradication, persistence and bistability. In case of a strong Allee effect (0 *< a <* 1), the only two possible regimes are eradication and bistability.

The condition for eradication is the same in all four models if we set *a* = *™*1. However as *a* grows, i.e when the Allee effect gets stronger, the ranges of *s* (fitness disadvantage for drive) and *r* (intrinsic growth rate) leading to population eradication get larger in the models with Allee effect ℬ𝒜 and 𝒟𝒜 (Appendix D and G).

Similarly when *a* grows, the ranges of *s* and *r* leading to population persistence get smaller in Models ℬ𝒜 and 𝒟𝒜 (Appendix E and G). We observe that the larger *a* is, the more the “persistence” and “bistable” regimes are restricted to high values of *r* and small values of *s* (Appendix G). In the case of a strong Allee effect (0 *< a <* 1), the “persistence” regime even disappears (Appendix E).

If the drive persists at the end, i.e. if there is a drive invasion or a coexistence state with a non-zero final population density, then, given how our models are formulated, the stronger the Allee effect, the smaller the final population size in Models ℬ𝒜 and 𝒟𝒜 (Appendix F.2).

#### 3.2.2 A density-dependence constraint on survival instead of fecundity might lead to a higher chance of eradication and reduces the final allelic density in case of drive persistence

The conditions for eradication or persistence are the same in Models ℬ𝒩 and 𝒟𝒩, i.e., they do not depend on whether the density dependence acts on births or deaths (no “bistable” regime for these models, Table 3 and Figure 2). However if we consider the models with Allee effect for a given *a* value, there is a higher chance of eradication when density dependence acts on deaths (Model 𝒟𝒜) than when it acts on births (Model ℬ𝒜) (Appendix D). Interestingly, when *r* → ∞, eradication is still possible in Model 𝒟𝒜 (for 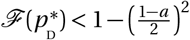) while it is not in Model ℬ𝒜 (Figure 2).

If the drive persists in models without Allee effect, i.e. if there is a drive invasion or a coexistence state with a non-zero final population density, the final population size is lower when the density-dependence constraint acts on the death rate (Model 𝒟𝒩) than on the birth rate (Model ℬ𝒩, see Appendix F.1). The same conclusion holds for models with Allee effect: for a given *a* value, if the drive persists, the final population is less dense in Model 𝒟𝒜 than in Model ℬ𝒜 (Appendix F.2).

### 3.3 A density-dependence constraint on the deaths instead of the births results in a faster invasion

We now focus on the speed of a drive invasion, i.e. the speed of the traveling wave emerging from a drive invasion (see Section 2.2).

A speed *v* of the wave can be calculated when the models are simplified (linearised) assuming low drive density. This speed corresponds to the speed of drive invasion when the movement of individuals is caused by the few drive individuals at the expansion edge, where the drive density is low. In this case, the wave is called a “pulled wave”. This happens when such small populations have high growth rates, because the movement is then mainly driven by new births. When movement is brought about by individuals in the bulk of the wave (i.e., in the case of a “pushed wave”), the calculated speed corresponds to a lower estimate of the speed: the real speed is higher, but cannot be calculated in general. In a previous article [28], we showed that the calculated speed *v* corresponds to the speed of a drive invasion when the dominance parameter *h* is lower than 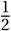, or for a drive fitness cost *s* small enough when 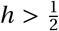 (for a precise condition, see [28]). This result was rigorously proven for large and small values of the intrinsic growth rate *r*, and numerically observed for all *r*. We calculate and compare this speed value in our four models (mathematical details are given in Appendix C.5).

In models ℬ𝒩 and ℬ𝒜 with density dependence acting on the birth term, this speed is given by:

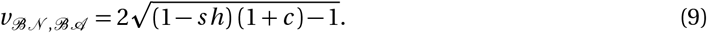

In models 𝒟𝒩 and 𝒟𝒜 with density dependence acting on the death term, the speed becomes:

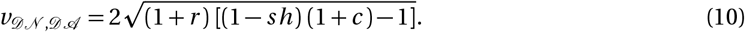

The speeds only exist for (1 *™ sh*)(1 + *c*) *>* 1 (or equivalently *s < s*_2_, with *s*_2,*g*_ given in equation (5)), which is the necessary condition to have a strictly positive drive allele production at the front of the wave. To understand why, first note that the density of drive alleles is very low at the front of the wave. Therefore, we can make the approximation that at least one parent in each couple formed at the front of the wave is a wild-type homozygote WW. Consequently, the offspring carrying a drive allele are necessarily heterozygotes: in the front of the wave, the production of drive alleles only relies on the heterozygotes. These heterozygotes have a fitness of (1 *™ sh*) and produce drive alleles at rate (1 + *c*): therefore, for a drive invasion to be possible, the production rate (1 *™ sh*)(1+*c*) of drive alleles should be above the rate 1 at which they disappear. The higher the production rate is, the faster the wave moves.

The speeds *v*_ℬ𝒩, ℬ𝒜_ and *v*_𝒟𝒩, 𝒟𝒜_ are very similar but differ by one coefficient: *v*_𝒟𝒩, 𝒟𝒜_ is 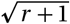 times larger than *v*_ℬ𝒩,ℬ 𝒜_. This difference relies on the density-dependence constraint, affecting either the births or the deaths. At the front of the wave, the population density, composed nearly only of wild-type individuals, reaches the maximum carrying capacity. Consequently, the density-dependence constraint hinders any increase in the population density and this happens in two different ways: in models ℬ𝒩 and ℬ𝒜, it limits the births so that they do not exceed the deaths, whereas in models 𝒟𝒩 and 𝒟𝒜, it increases the death rate to compensate the births. As a result, the turnover rate is greater in models 𝒟𝒩 and 𝒟𝒜, which induces a faster invasion because the wave movement is mainly driven by new births. Details of the speed calculations are given in Appendix C.5. To illustrate this result, we plot the speed of the wave for the four models in Figure 3 and observe that the speed of the drive invasion always increases with *r* in models 𝒟𝒩 and 𝒟𝒜, in contrast with models ℬ𝒩 and ℬ𝒜, for which the speed of the wave does not depend on *r*. Note however that while the speed *v*_ℬ𝒩, ℬ𝒜_ is independent of *r* for a drive fitness cost *s* small enough, it is not the case for the final density 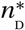, which depends on *r* (Table 3 and Figure 2, models ℬ𝒩 and ℬ𝒜). As a result, for a small enough *s* in models and, the wave travels at a constant speed no matter the density of population left behind.

### 3.4 The Allee effect might cause the failure of threshold dependent drives

Finally, in Figure 3, we observe that for *s > s*_2,*g*_, the Allee effect might prevent drive invasion. Yelloworange areas in heatmaps (a) (resp. (d)) are becoming blue in heatmaps (b) and (c) (resp. (e) and (f)) due to the Allee effect in the model. Noticeably, the drive invasions for *s > s*_2,*g*_ are thresholddependent drive invasions, often considered as more socially responsible than threshold independent drives invasions [25, 26]. Once again this influence is accentuated when the Allee effect gets stronger (for larger values of *a*). Allee effects may therefore hamper the spread of threshold-dependent drives.

**Figure 3:**
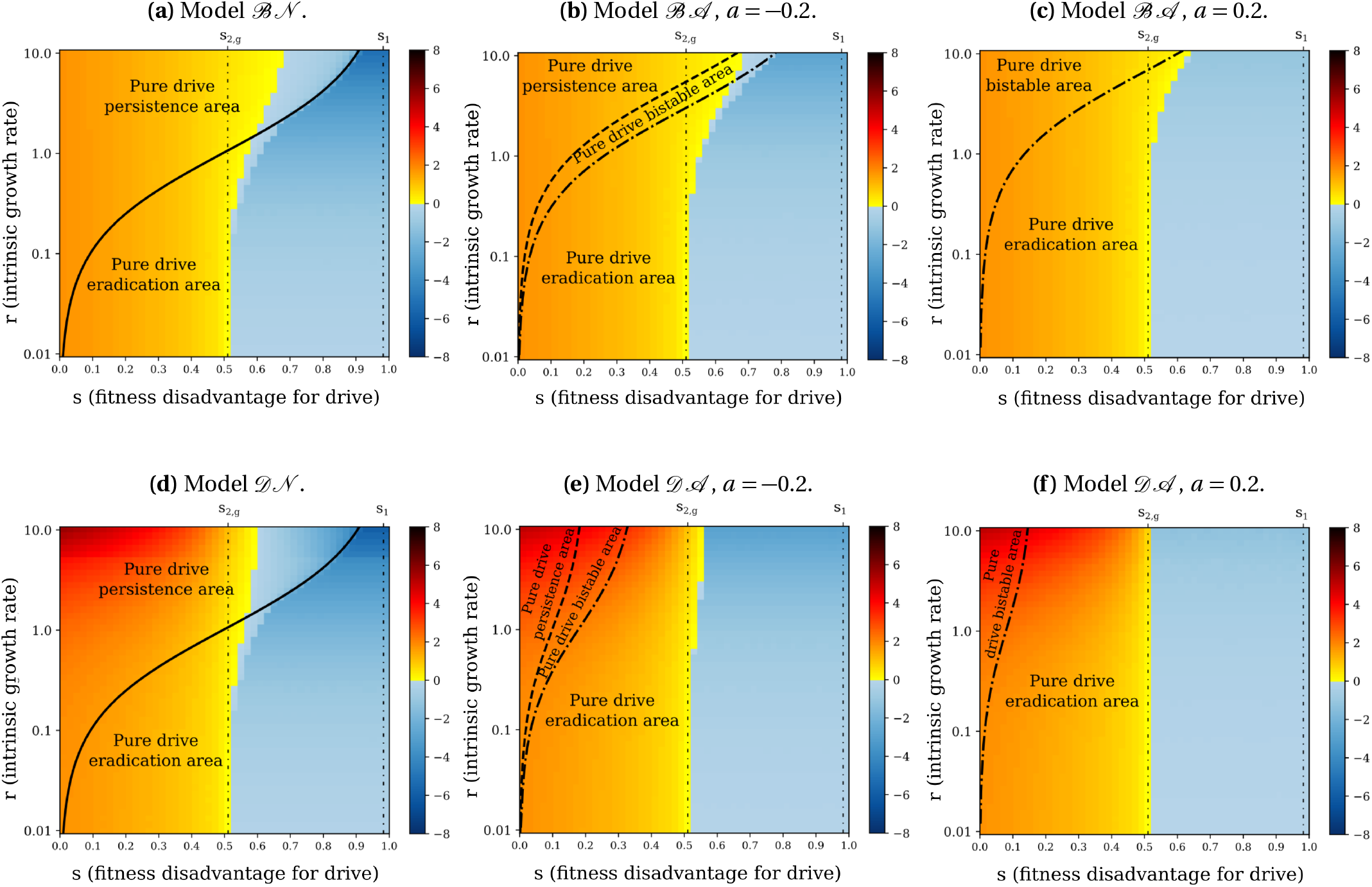
Speed of the wave of drive alleles shown as shades of color, with parameters *c* = 0.85 and *h* = 0.9. When the drive invades the population, the speed is positive (in yellow-orange). On the contrary, when the wild-type invades the population, the speed is negative (in blue). The “pure drive” areas correspond to the final population density expected in case of a drive invasion: persistence (*n*^*^ = *n* ^+^), eradication (*n*^*^ = 0) or bistability (either *n*^*^ = *n* ^+^ or *n*^*^ = 0 depending on the initial condition). Since 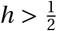, coexistence of drive and wild-type alleles in the final population is impossible (see Table 2b). Each model is initiated with the initial conditions described in Figure S1. For *s*_2,*g*_ *< s < s*_1_, the speed value may vary if we consider different initial conditions (bistability on the final allele frequencies, see Table 2b). On the left of the vertical line *s* = *s*_2,*g*_, the speed value is independent of *r* when the density-dependence constraint acts on fecundity (Models ℬ𝒩 and ℬ𝒜) while it grows with *r* when the density-dependence constraint acts on survival (Models 𝒟𝒩 and 𝒟𝒜).

## 4 Discussion

Understanding the conditions for the spatial spread of an artificial gene drive and its consequences on a targeted population is essential before considering any field release. Laboratories experiments provide information on gene drive dynamics in a small confined and controlled environment, and mathematical models can help gain further insights at small and larger scales.

Theoretical models are meant to provide insights on real-world dynamics, so it is important to assess how the result of a model depends on modelling choices. In this article, we investigated the influence of considering i) demography, and more precisely different values of the intrinsic growth rate of the target population, ii) the presence/absence of an Allee effect and iii) which fitness component (birth or death) is affected by density dependence. We considered the effects of these features on the type of outcome, on final population density, and on the speed of the wave.

We first described the different qualitative outcomes, extending results from our previous studies [11, 28] on the importance of taking into account demography in the models. We confirm that the intrinsic growth rate *r* qualitatively affects results at intermediate values of the fitness cost *s*. A high intrinsic growth rate leads to a threshold-dependent drive invasion, while a low intrinsic growth rate results in the decay of drive alleles uniformly in space. Models not considering population densities but focusing on frequencies [e.g. 18, 22–25] have dynamics similar to our models provided *r → ∞*.

As intuitively expected, an Allee effect makes the population more susceptible to eradication, widening the range of *s* (fitness disadvantage for drive) and *r* (intrinsic growth rate) values leading to population eradication after a drive invasion. This phenomenon is accentuated when the Allee effect gets stronger (for larger values of *a*). In addition, given the way our models are formulated, in models with Allee effect, the larger *a*, the lower the final population density in case of drive persistence, meaning that an Allee effect might represent a non-negligible helping force to eradicate or suppress natural populations. However, we also showed that an Allee effect might reduce the range of *s* and *r* values leading to a threshold-dependent drive invasion, often considered as more socially responsible than threshold independent drives invasions [25, 26].

Finally, we considered the impact of whether the density-dependence constraint targets births or deaths: close to the maximal carrying capacity, in case of rarefaction of the resources, the net growth of the population is limited by either a low number of offspring per generation or a high death rate.

In this study, we show that when density dependence acts on deaths, it acts in concert with the Allee effect by enlarging the eradication conditions and reducing the final density, compared to when density dependence acts on births. How density dependence acts also strongly impacts the speed of propagation: a drive invasion would be 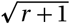 times faster for a density-dependence constraint on the death rate instead of the birth rate. This prediction holds for a fitness cost reducing the birth rate (individuals carrying drive alleles have fewer offspring than wild-type ones). However, the conclusions might change for a fitness cost increasing the death rate instead, as shown in a different model of CRISPR-based homing drives [31].

Our models are deterministic. This deterministic framework can describe population dynamics at large scales, but cannot capture “chasing” events, whereby wild-type individuals recolonize empty space in the wake of the wave of an eradication drive, and which can arise at low population densities [43–47]. Stochastic fluctuations are likely to be important in particular in the case of suppression and eradication drives, and are left for future investigation.

Among the deterministic models in the literature, the models we develop are generalist: they could be applied to different species, and any gene drive construct reducing the fitness of the individual carrying it. These models do not aim to bring precise and quantitative predictions, for which more specific models need to be developed, but rather get some insights into the possible outcomes, and dissect the roles played by different model elements. However, this generalist approach naturally comes with simplifications.

In our models, we assume that gene conversion either successfully takes places, or does not take place. We did not include resistance alleles which can emerge when conversion fails and repairs by non-homologous end-joining occur, or resistance due to standing genetic variation at the target locus. The emergence of resistance alleles can alter the propagation of the drive, but can be mitigated by specific constructs [18, 48–50].

Some other simplifications are directly related to the biological characteristics of the species. The polyandrous mating system of mice populations can limit the spread of gene drives [51, 52] or mate search capabilities [53]. In mosquito populations, the plural life stages (egg, larva, pupa and adults) might influence the modelling conclusions and need to be taken into account by including corresponding age structure in models [54–56]. In bee populations, the haploid phases of the life cycle result in less powerful drives: the conditions for fixation are narrower and the spread is slower [57, 58]. Finally, it is not rare that males and females have different fitnesses in transgenic mosquitoes [8, 16, 47, 59]: more specific models than ours would need to include sex differences.

Finally and more broadly, species do not live in isolation, and interactions of the targeted species within its ecosystem would need to be considered. Competing species or predators can facilitate drive-based suppression [57], and environmental conditions such as seasonality (dry or wet season) can highly impact the eradication of mosquito populations, for example [44, 46, 47]. It is of public utility to also consider the impact of gene drive on the whole ecosystem and anticipate the potential risks: the probability of transmission of the gene drive cassette to another species [60], or the cascade of population dynamics and evolutionary processes potentially initiated by the eradication of a species [61].

Overall, we have shown the importance of considering precise population dynamics on the outcome of the release of a drive. This approach through theoretical models gives first interesting insights that now need to be enhanced with ecological knowledge on specific systems.

## Appendix

### A Initial conditions

All our simulations in one-dimensional space are initiated with the initial conditions described in the main text, and illustrated in Figure S1:

**Figure S1:**
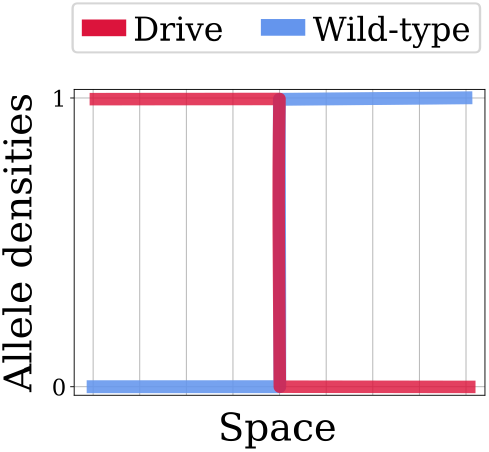
Initial conditions used in the simulations. The left half of the domain is full of drive (*n*_DD_ = *n*_*D*_ = 1), and the right half is full of wild-type (*n*_WW_ = *n*_*W*_ = 1).

### B Allee effect

We consider the equation describing the dynamics of the population density *n* :

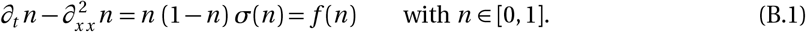

The Allee effect characterises a positive correlation between population density and the per capita population growth rate.

Without Allee effect, the population growth rate *f* (*n*) is always positive and the per capita population growth rate 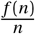 is maximum as the population density *n* tends to zero. This happens for example when *σ*(*n*) = 1 in Equation (B.1) (Figure S2a). Mathematically, we write:

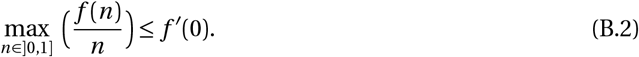

With a weak Allee effect, the population growth rate *f* (*n*) is still positive, but the maximum of the per capita population growth rate 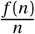 is reached at a strictly positive population density. This happens for example when *σ*(*n*) = (*n* − *a*) with −1 *< a <* 0 in Equation (B.1) (Figure S2b). Mathematically:

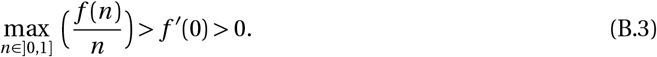

Finally, with a strong Allee effect, the population growth rate *f* (*n*) is negative for small population density, and positive after. This happens for example when *σ*(*n*) = (*n ™ a*) with 0 *< a <* 1 in Equation (B.1) (Figure S2c). Mathematically:

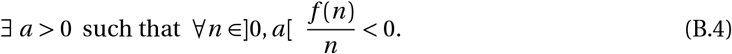

**Figure S2:**
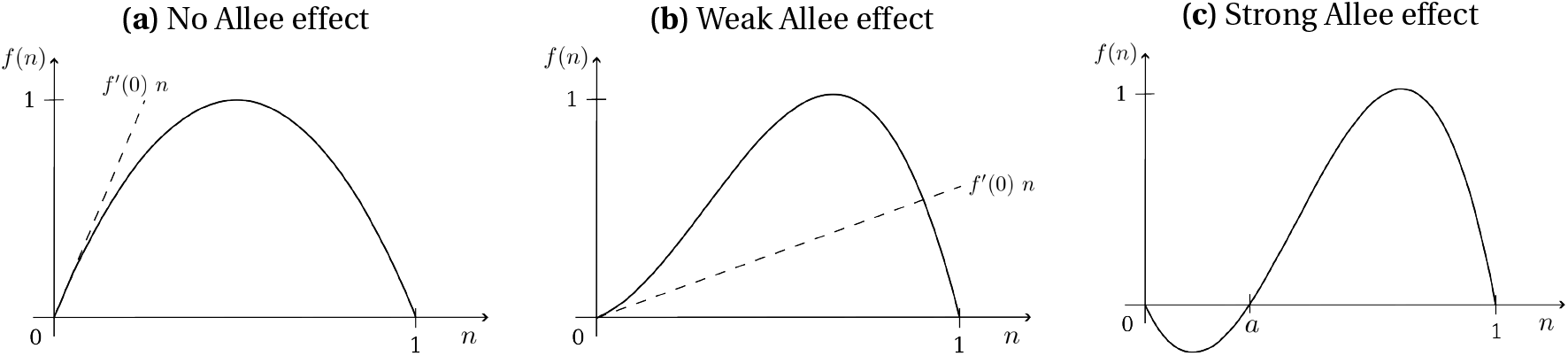
Illustration of the three cases concerning the Allee effect.

### C Models

#### C.1 Genotype densities

For the sake of clarity, we omit variables in the notation (*n*_*i*_ = *n*_*i*_(*t, x*)) in the following. Each model contains three equations for the three genotype densities: homozygote drive *n*_DD_, heterozygote *n*_DW_ and homozygote wild-type *n*_WW_.

Model ℬ𝒩:

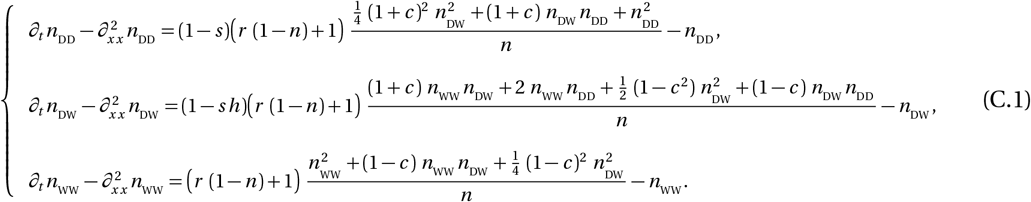

Model ℬ𝒜:

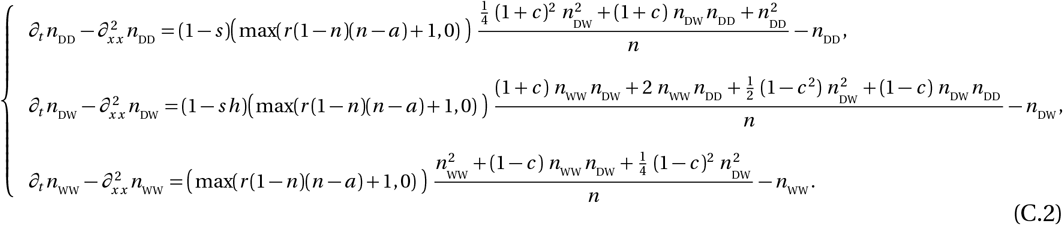

Model 𝒟𝒩:

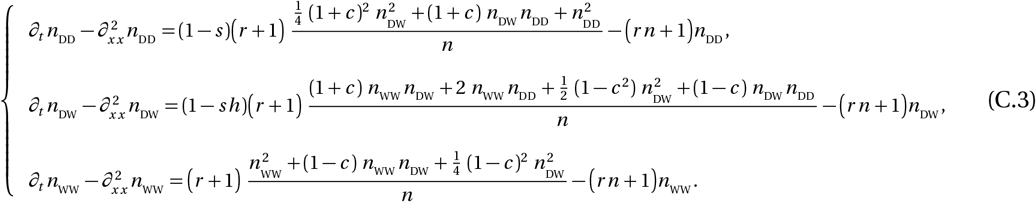

Model 𝒟𝒜:

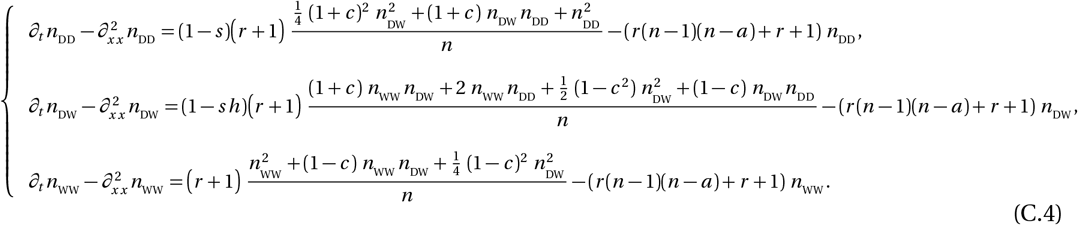

Note that all four models reduce to a single model for *r* = 0. This model is:

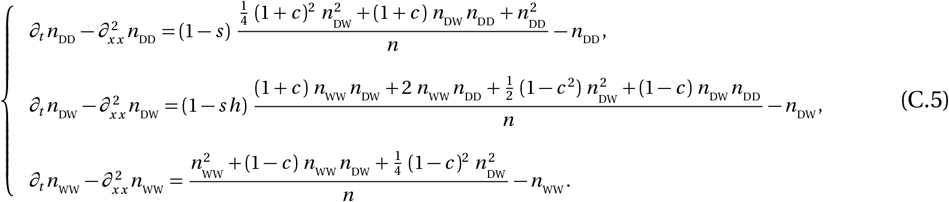

#### C.2 Allelic densities

For our analysis, it is convenient to introduce the allelic (half-) densities (*n*_D_, *n*_W_). For a conversion occurring in the germline, we have 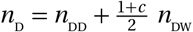 and 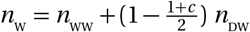 (see section 3.2 in [28] for more details). We deduce the following systems.

Model ℬ𝒩:

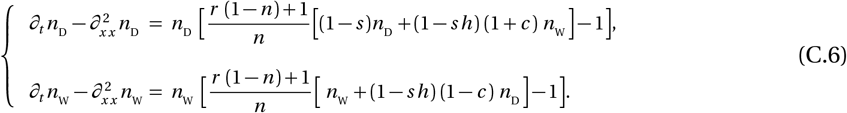

Model ℬ𝒜:

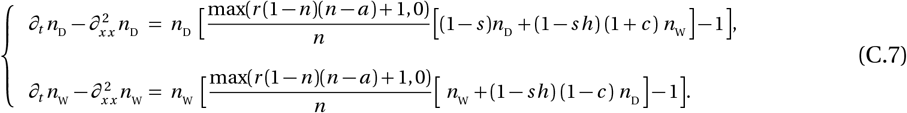

Model 𝒟𝒩:

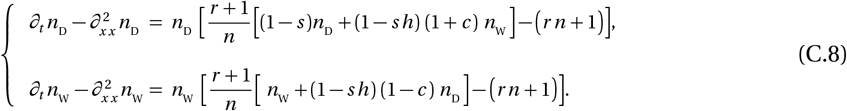

Model 𝒟𝒜:

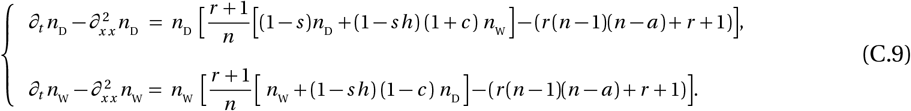

#### C.3 Allelic frequencies

It may sometimes be more appropriate to write the models in terms of the proportion of drive allele 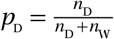 and total population size *n* = *n*_D_ + *n*_W_. The models become the following.

Model ℬ𝒩:

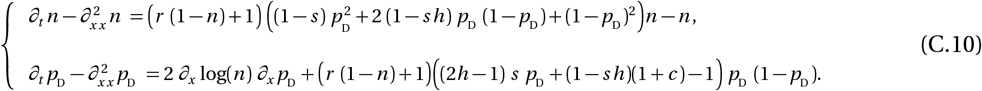

Model ℬ𝒜:

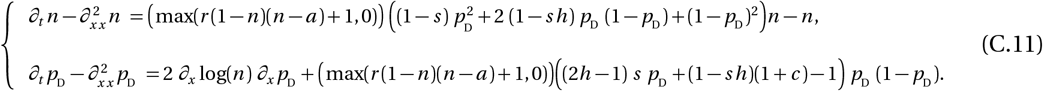

Model 𝒟𝒩:

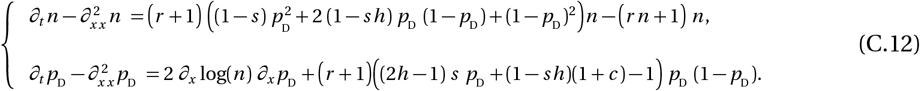

Model 𝒟𝒜:

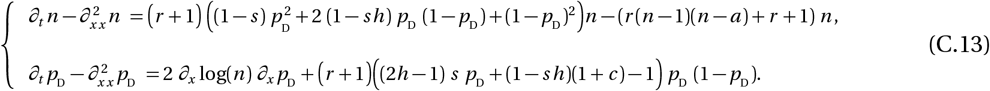

Equations on *p*_D_ differ from the standard equation often used in populations genetics, as they contain an advection term 2 *∂*_*x*_ (log *n*) *∂*_*x*_ *p*_D_. This term appears when calculating 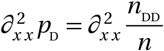 and represents a demographic flux from denser to less dense areas, due to variations in population density. It is opposed to the spread of the costly drive allele (see Figure 2 [11]).

#### C.4 Final allelic proportions for *r* small and large

In models ℬ𝒩 and ℬ𝒜 for large values of *r*, using the Strugarek-Vauchelet rescaling [62] in (C.10) and (C.11), the systems reduce to one limit equation on *p*_D_ :

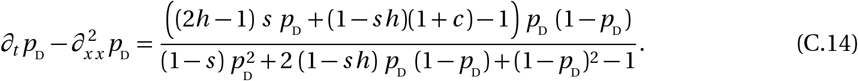

Equation (C.14) has already been studied in [18]: a heatmap illustrates the final proportions in the case *c* = 0.85 (Figure 4 in [18]). In models 𝒟𝒩 and 𝒟𝒜, the equation on *p*_D_ in (C.10) and (C.11) is:

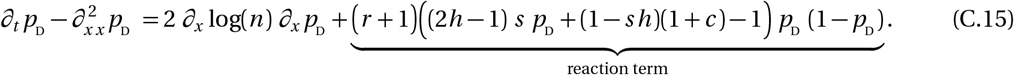

The reaction term in equation (C.15) becomes larger as *r* increases: this indicates that the traveling wave has an infinite speed when *r* tends to infinity, meaning that the equilibrium is reached instantaneously. Therefore, the term 2 *∂*_*x*_log(*n*) *∂*_*x*_ *p*_D_ is instantaneously zero and the final proportions are the same as for models ℬ𝒩 and ℬ𝒜.

As a consequence, all models ℬ𝒩, ℬ𝒜, 𝒟𝒩 and 𝒟𝒜 share the same final proportions for large values of *r*. This conclusion also holds for *r* = 0, as the models are equal (see Appendix C.1). These proportions have already been determined in a previous article [28], for the ℬ𝒩 case. We recall these results below in section F and generalise them to our four models.

#### C.5 Speed of the problem simplified at low drive density

In Section 3.3, we focus on drive invasion and therefore consider low drive density and high wildtype density at the front of the wave. The speed *v* of the wave can be calculated when the models are simplified (linearised) at low drive density: it is deduced from the reproduction of the few drive individuals at the front wave,

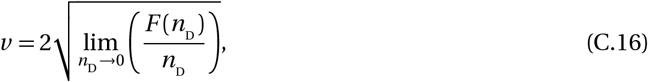

where *F* represents the net production of drive alleles. The formulas for *F* in the different models are the following.

Model ℬ𝒩:

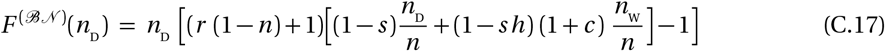

Model ℬ𝒜:

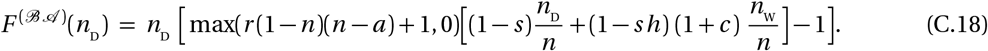

Model 𝒟𝒩:

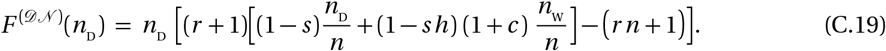

Model 𝒟𝒜:

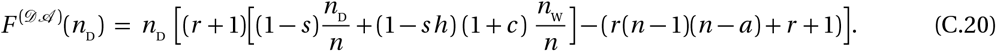

Considering high wild-type density at the front of the wave (*n*_W_ ≈ *n* ≈ 1), the speed in models ℬ𝒩 and ℬ𝒜 is given by:

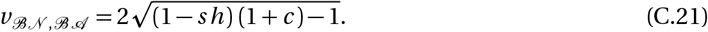

In models 𝒟𝒩 and 𝒟𝒜, we have:

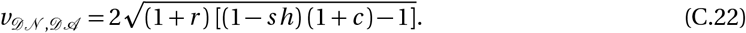

To understand why *v*_𝒟𝒩, 𝒟𝒜_ is greater by a coefficient of 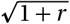 than *v*_ℬ𝒩, ℬ𝒜_, we have to understand the population dynamics at the front of the wave. There, the density is close to the maximum carrying capacity 1, with low drive density (*n*_D_ *≈*0) and high wild-type density (*n*_W_ *≈ n ≈* 1). On one hand, in models ℬ𝒩 and ℬ𝒜, the density-dependence constraint is placed on the birth term, reducing the production rate of drive alleles to (1 *™ sh*) (1 + *c*) while individuals disappear at rate 1 (C.17, C.18). On the other hand, in models and, the density-dependence constraint is placed on the death term increasing to (*r* + 1) the rate at which the drive alleles disappears, while they are produced at rate (*r* + 1)(1 − *sh*)(1 + *c*) (C.19, C.20). Consequently, the net production remains constant (1 − *sh*)(1 + *c*), but the turnover rate is *r* + 1 times greater. As the wave movement largely relies on the reproduction, this reflects in the speed formula: the propagation is 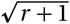 times faster.

### D Comparison of the conditions leading to eradication

To compare the conditions leading to eradication, we refer to the results summarized in Table 3.

In Models ℬ𝒩 and 𝒟𝒩, eradication occurs when:

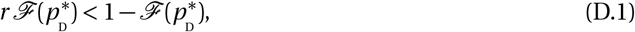

in Model ℬ𝒜, when:

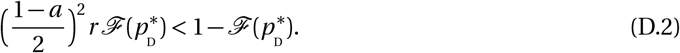

and in Model 𝒟𝒜, when:

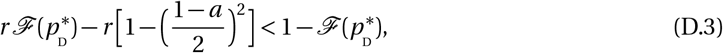

With 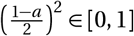 and 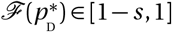, we obtain the following inequalities:

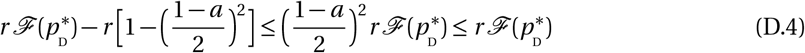

Inequalities (D.2) and (D.3) both imply (D.1). In other words, eradication is easier to achieve in Models ℬ𝒜 and 𝒟𝒜 (with Allee effect) than in Models ℬ𝒩 and 𝒟𝒩 (without Allee effect) in the sens that the range of parameters leading to eradication is larger.

Inequality (D.3) also implies (D.2), i.e., in models with Allee effect, for a given *a* value, the density-dependence constraint on the deaths (Model 𝒟𝒜) makes the eradication easier than if it is on the births (Model ℬ𝒜).

Note that when *a* = *™* 1, the conditions leading to eradication are equivalent in all four models. When *a* becomes larger, the range of parameters leading to eradication in models ℬ𝒜 and 𝒟𝒜 also becomes wider.

### E Comparison of the conditions leading to persistence

To compare the conditions leading to persistence, we refer to the results summarized in Table 3.

In Models ℬ𝒩 and 𝒟𝒩, persistence occurs when:

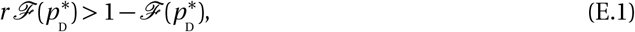

in Model ℬ𝒜, when:

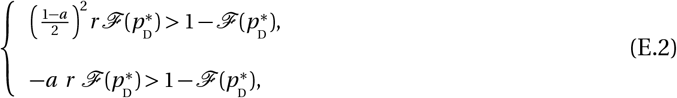

and in Model 𝒟𝒜, when:

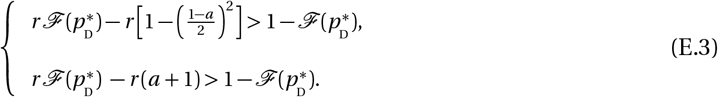

First note that persistence is only possible in case of a weak Allee effect (*a <* 0) in Models ℬ𝒜 and 𝒟𝒜 (second lines in Systems (E.2) and (E.3)). With 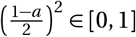 and 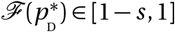, we obtain the following inequalities:

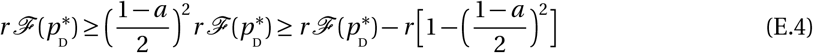

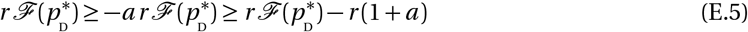

Inequality (E.1) imply both (E.2) and (E.3). In other words, persistence is easier to achieve in Models ℬ𝒩 and 𝒟𝒩 (without Allee effect) than in Models and (with Allee effect) in the sens that the range of parameters leading to persistence is larger.

Inequality (E.2) also implies (E.3), i.e., in models with Allee effect, for a given *a* value, the density-dependence constraint on the births (Model ℬ𝒜) makes the persistence easier than if it is on the deaths (Model 𝒟𝒜).

Note that when *a* = −1, the conditions leading to persistence are equivalent in all four models. When *a* becomes larger, the range of parameters leading to persistence in models ℬ𝒜 and 𝒟𝒜 becomes smaller. In case of a strong Allee effect (*a >* 0), the “persistence” regime disappears.

### F Comparison of the final densities in case of persistence

#### F.1 Comparison of the final density in models ℬ𝒩 and 𝒟𝒩

We compare the final densities for models ℬ𝒩 and 𝒟𝒩 given in Table 3 in case of persistence.

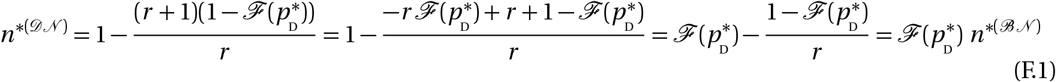

With 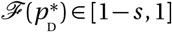. The final density is 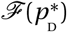 times lower in model 𝒟𝒩 than in model ℬ𝒩, in case of persistence. Note that 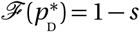 in case of a drive invasion, and 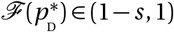 in case of coexistence.

#### F.2 Comparison of the final density in models ℬ𝒜 and 𝒟𝒜

We compare the final densities for model ℬ𝒜 and 𝒟𝒜 in case of persistence given in Table 3.

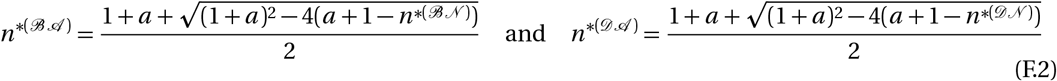

From the previous section F.1, we know that 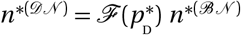 with 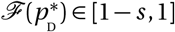. Therefore:

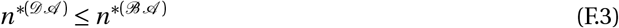

This inequality is strict in case of a drive invasion 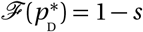 or in case of coexistence 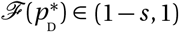. We derive both density as functions of *a*, in case of drive persistence 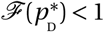:

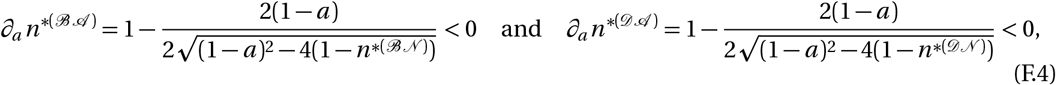

as *n*^*(ℬ𝒩)^ *<* 1 and *n*^*(𝒟𝒩)^ *<* 1 when 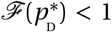 (see Table 3). In other words in case of a drive persistence, the stronger the Allee effect, the smaller the final population density in models ℬ𝒜 and 𝒟𝒜.

### G Final population density when the strength of the Allee effect varies, in case of drive invasion

The “pure drive” areas described in Figure 2 correspond to the final density expected in case of a drive invasion: persistence (*n*^*^ = *n* ^+^), eradication (*n*^*^ = 0) or bistability (either *n*^*^ = *n* ^+^ or *n*^*^ = 0 depending on the initial conditions). These final densities are given in Table 3 with 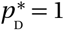 (the proportion of drive allele in the final population is one in case of a drive invasion without coexistence). Without Allee effect, there only exists pure drive eradication and persistence areas (Models ℬ𝒩 and 𝒟𝒩). However, if we consider an Allee effect (Models ℬ𝒜 and 𝒟𝒜), a pure drive bistable area appears. In Figure S3, we plot the boundary lines delimiting the pure drive areas in models with Allee effect, for different values of *a*. The first boundary line, delimiting the pure drive persistence area (above it) from the pure drive bistable area (under it), will be referred to as the *upper boundary line* and is represented with a dash-dotted line (· · -). The second boundary line delimiting the pure drive bistable area (above it) from the pure drive eradication area (under it), will be referred to as the *under boundary line* and drawn with a dashed line (-).

**Figure S3:**
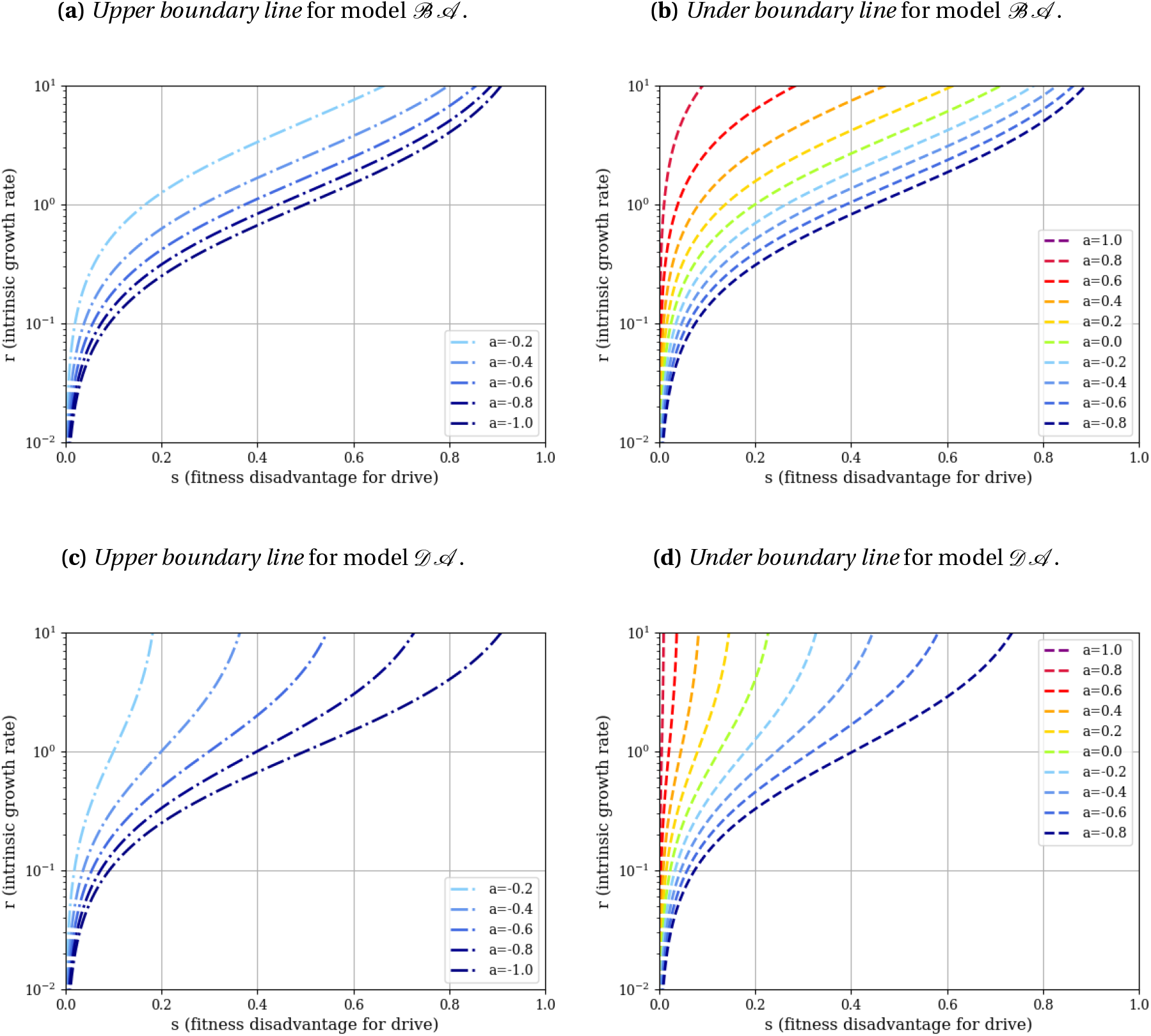
Boundary lines for different values of *a* in models with Allee effect ℬ𝒜 and 𝒟𝒜. The *upper boundary line* plotted on panels (a) and (c), delimits the pure drive persistence area (above it) from the pure drive bistable area (under it). It is represented with a dashdotted line. The *under boundary line* plotted on panels (b) and (d), delimits the pure drive bistable area (above it) from the pure drive eradication area (under it). It is represented with a dashed line. These boundaries were represented for two values of *a* in Figures 2 and 3 (*a* = −0.2 and *a* = 0.2).

The higher *a*, the larger the pure drive eradication area and the more the pure drive persistence and bistable areas are restricted to high values of *r* and small values of *s*. In case of a strong Allee effect (0 *< a <* 1), the pure drive persistence area even disappears; this is why only negative values of *a* are plotted in Figure S3 (a) and (c).

## Conflict of interest

The authors declare that they have no conflict of interest.

## Acknowledgements

This work is funded by ANR-19-CE45-0009-01 TheoGeneDrive. This project has received funding from the European Research Council (ERC) under the European Union’s Horizon 2020 research and innovation programme (grant agreement No 865711).

We are grateful to the INRAE MIGALE bioinformatics facility (MIGALE, INRAE, 2020. Migale bioinformatics Facility, doi: 10.15454/1.5572390655343293E12) for providing computing resources

## Footnotes

** The term “drive invasion” here is slightly ambiguous, as it does not specify the genetic composition in the wake of the eradication wave. This exponentially small population might contain the three possible genotypes or only the drive-homozygous one, depending on parameter choices.

## Notes

### Competing Interest Statement

The authors have declared no competing interest.

